# Visual evoked feedforward-feedback travelling waves organize neural activity across the cortical hierarchy in mice

**DOI:** 10.1101/2021.08.09.455716

**Authors:** Adeeti Aggarwal, Connor Brennan, Jennifer Luo, Helen Chung, Diego Contreras, Max B. Kelz, Alex Proekt

## Abstract

Sensory processing is distributed among many brain regions that interact via feedforward and feedback signaling. It has been hypothesized that neuronal oscillations mediating feedforward and feedback interactions organize into travelling waves. However, stimulus evoked travelling waves of sufficient spatial scale have never been demonstrated directly. Here, we show that simple visual stimuli reliably evoke two traveling waves with spatial wavelengths that cover much of the cerebral hemisphere in awake mice. 30-50Hz feedforward waves arise in primary visual cortex (V1) and propagate rostrally, while 3-6Hz feedback waves originate in the association cortex and flow caudally. The phase of the feedback wave modulates the amplitude of the feedforward wave and synchronizes firing between V1 and parietal cortex. Altogether, these results provide direct experimental evidence that visual evoked travelling waves percolate through the cerebral cortex and coordinate neuronal activity across broadly distributed networks mediating visual processing.

## Introduction

Feedforward and feedback signaling contribute to the hierarchical processing of sensory stimuli, creating predictions and attaching behavioral context to the sensory world (Felleman and Van Essen, 1991; Bosman et al., 2012; Bastos et al., 2015; van Ede et al., 2015; Mejias et al., 2016; Michalareas et al., 2016; Vecchia et al., 2020). Feedforward processing involves bottom-up assembly of abstract stimulus representations in higher-order areas from simple receptive fields in the primary cortex (Felleman and Van Essen, 1991; Markov et al., 2014). Feedback processing, in contrast, involves top-down influences such as attention, prediction, and context (Felleman and Van Essen, 1991; Markov et al., 2014). Formulating predictions about the next sensory stimulus or deciding which stimulus to pay attention to requires temporal integration (Friston, 2008; Bastos et al., 2015; Friston and Buzsáki, 2016; Posner et al., 2018). Thus, it is thought that feedback modulation evolves on a slower time scale relative to feedforward processing (Bosman et al., 2012; Bastos et al., 2015; Alamia and VanRullen, 2018).

Feedforward – feedback interactions between the different cortical regions involved in sensory processing must be coordinated to give rise to integrated percepts situated in the behavioral context. The role of neuronal oscillations in coordinating neuronal activity has been a subject of intense investigation, especially in primate vision. By analyzing individual pairwise interactions between neural oscillations present at different areas of the primate cortex, many prominent studies have shown that feedforward processing involves gamma oscillations, whereas feedback signaling uses alpha (8-12Hz) oscillations (Engel et al., 2001; Bosman et al., 2012; Van Kerkoerle et al., 2014; Bastos et al., 2015; Michalareas et al., 2016). Thus, consistent with their presumed behavioral roles, feedback signalling utilized slower temporal oscillations compared to feedforward channels.

Pairwise interactions between oscillations in different cortical sites during processing of sensory stimuli raise several fundamental questions. Do pairwise feedforward and feedback interactions give rise to a single coherent assembly that coordinates activity among the different cortical regions involved in processing sensory stimuli? How does the brain coordinate the feedforward and feedback processing given the significant differences in timescales? One possibility for a neurophysiological process that could coordinate activity amongst multiple regions in the processing hierarchy is a spatiotemporal travelling wave. Early EEG work identified travelling waves in the feedforward and feedback directions (Adrian and Matthews, 1934; Goldman et al., 1948; Adrian and Yamagiwa, 1960; Darrow and Hicks, 1965; Hughes, 1995). However, due to the low spatial resolution of the EEG, the interpretation of these findings is unclear. Indeed, travelling waves recorded directly from the cortical surface have different speeds and propagation patterns compared to their EEG counterparts (Hangya et al., 2011; Bahramisharif et al., 2013; Mak-McCully et al., 2015; Muller et al., 2018). Both spontaneous and stimulus evoked travelling wave-like phenomena have been identified using voltage sensitive dyes and neurophysiological recordings from brain parenchyma in the primary and higher order visual areas (Bringuier et al., 1999; Gabriel and Eckhorn, 2003; Benucci et al., 2007; Xu et al., 2007a; Nauhaus et al., 2009; Polack and Contreras, 2012; Sato et al., 2012; Muller et al., 2014; Besserve et al., 2015; Townsend et al., 2015; Davis et al., 2020). Most of this work however, focused on a single cortical area rather than inter-area communication. Some studies attempted to identify spatiotemporal waves that span multiple cortical sites and concluded that sensory stimuli trigger two independent cortical waves, which travel along the horizontal fiber network in each site (Polack and Contreras, 2012; Muller et al., 2014). Other studies identified a reflective boundary between the primary and the secondary visual cortex (Xu et al., 2007b). Thus, while a single spatiotemporal wave of activity offers an attractive possibility for coordinating cortical activity, the existence of stimulus-evoked travelling waves with sufficient spatial scale to span the cortical hierarchy has never been directly demonstrated. Furthermore, the relationship between feedforward-feedback processing of sensory stimuli and the travelling waves evoked by them in the cortex has not been clarified.

We deploy a combination of high-density neurophysiological recordings and analytic techniques to identify large scale spatiotemporal patterns of neuronal activity evoked by a single presentation of a simple, supra-threshold visual stimulus. By focusing our analyses on global activity patterns, rather than pairwise interactions, we show that both the feedforward and feedback aspects of visual evoked activity form travelling waves that percolate through much of the cortex in awake mice. Feedforward waves have a fast (30-50Hz) temporal frequency and propagate from V1 rostrally. Feedback waves are characterized by a slow (3-6Hz) oscillation, thought to be a rodent analogue of the primate alpha oscillation. These feedback waves propagate caudally from association cortices towards V1. The phase of the feedback wave modulates the amplitude of the feedforward wave, thereby forming a single multiplexed visual evoked spatiotemporal response. Finally, we demonstrate that the feedback wave entrains firing of individual neurons in both V1 and in parietal association cortex. As a consequence, following stimulus presentation, previously uncorrelated firing in V1 and parietal cortex phase lock their firing in relation to the stimulus to form a transient neuronal assembly. Thus, we provide direct evidence that feedforward-feedback interactions organize into large scale traveling waves evoked by simple visual stimuli. These waves serve as a scaffold that coordinates neural firing across distant cortical areas.

## Results

Our primary goal is to experimentally define salient spatiotemporal signatures of responses to simple visual stimuli. To accomplish this, we performed high density *in vivo* electrophysiological recordings in awake head fixed mice (n = 13) (Methods for verification of wakeful states, Sup. Figure 1, Sup. Video 1). Local field potentials (LFPs) were recorded from the dural surface using a 64 channel electrocorticography (ECoG) grid placed over the left hemisphere (Figure 1a). Two 32 channel laminar probes were also inserted perpendicular to the cortical surface targeting the primary visual cortex (V1) and the posterior parietal area (PPA). Histological and neurophysiological (Methods) localizations of the laminar probes were used to triangulate the stereotaxic locations of the individual ECoG electrodes. The ECoG grid covered a significant fraction of the cerebral hemisphere including visual, association, retrosplenial, somatosensory, and motor/frontal areas (Figure 1b).

**Figure 1:**
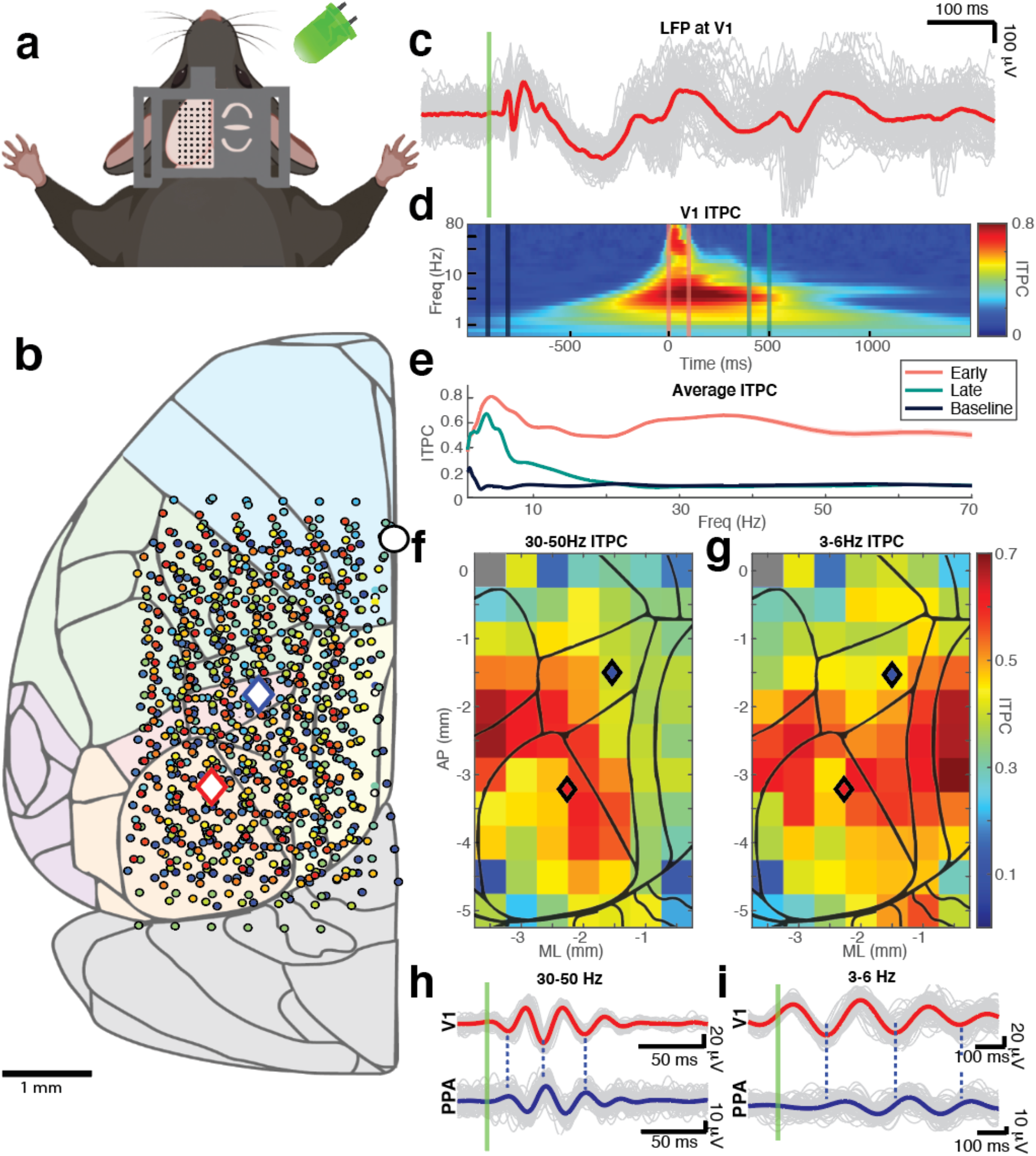
Visual stimuli elicit strong intertrial phase coherence over large cortical areas. a. Schematic showing the 64 channel electrocorticography (ECoG) grid used to record local field potentials (LFPs) from the cortical surface of the left hemisphere of 13 awake mice. Stimuli consisted of 10 ms flashes of a green LED placed in front of the R eye (100 trials, intertrial interval 3-4s). b. Stereotaxic coordinates of ECoG electrodes from 13 animals (color coded by animal). V1 and PPA targets for laminar probes are shown by red and blue diamond respectively. The white circle marks bregma. The cortical surface is shaded by area according to the Allen Brain Atlas (Franklin, Keith, B. and Paxinos, 2007; Lein et al., 2007; Reference Atlas :: Allen Brain Atlas: Mouse Brain): visual (orange), association (red), retrosplenial (yellow), somatosensory (green), motor/frontal (blue), and cerebellum (gray). c. Single trial and average visual evoked potentials (VEPs) over V1 are indicated by gray and red lines respectively. Stimulus onset is denoted by the green line. d. Inter-trial phase coherence (ITPC) computed at V1 and averaged over single trials and animals (0 ms marks stimulus onset). e. Time slices through the coherogram (Figure 1D). Colors of the traces correspond to time segments shown in D. Thick line shows the average. Shaded areas show 95% confidence intervals. f. Average ITPC of 30-50Hz oscillations within the first 100 ms of the VEP averaged over animals at each stereotaxic location. Locations in which ITPC does not meet Bonferroni corrected statistical significance compared to time shuffled surrogate data are shaded in gray. g. Similar to E for average ITPC of 3-6Hz activity within the first over 800 ms of the VEP. h. Top: VEPs recorded over V1 and filtered at fast (30-50 Hz) oscillations (gray and red show single trials and trial average respectively). Bottom: Same data recorded from over PPA (gray and blue show single trials and trial average). Green line shows stimulus onset. Dashed lines highlight the phase offset between V1 and PPA. i. Similar to G except the signals are filtered at 3-6Hz. Slower (3-6Hz) oscillations also show a phase shift between V1 and the PPA. *Data in C, H, and I are from a single representative mouse.

### Simple, brief visual stimuli evoke widespread time-locked coherent oscillations at both high (30-50Hz) and low (3-6Hz) frequencies

As in previous work (Childers et al., 1987; Liberati et al., 1991; Churchland et al., 2010; Aggarwal et al., 2019), the visual-evoked potential (VEP) in V1 (Methods) varies from trial-to-trial. Nevertheless, early fast oscillations (30-50Hz) followed by longer lasting slow oscillations (3-6Hz) are reliably identified from trial to trial (Figure 1c). Analysis of the inter-trial phase coherence (ITPC) confirms that these two oscillations are consistently phase locked to the stimulus (Stouffer’s *p-values*<0.00001 compared to time shuffled data) (Figure 1d). ITPC computed over the first 100 ms after the stimulus reveals two peaks centered at 3-6Hz and 30-50Hz. The 3-6Hz oscillation remains coherent for 500ms after the stimulus (Figure 1e). Because these two oscillations (fast, 30-50Hz and slow, 3-6Hz) are reliably phase locked to the stimulus in all mice, we focus our subsequent analyses on these oscillations.

Phase locking of fast and slow oscillations to the stimulus is not limited to V1. Both fast and slow oscillations are phase locked to the stimulus across much of the cortical surface (Figure 1f and 1g, respectively). Phase locking to the stimulus over large areas of the cortical surface strongly suggests that oscillations recorded at different sites are interdependent. Consistent with this suggestion, LFPs filtered at fast and slow frequencies in V1 and PPA exhibit phase coupling (Figure 1h and 1i). Interestingly, oscillations at both temporal frequencies have a non-zero phase lag between V1 and PPA (Figure 1h, and Figure 1i). This raises the possibility that the stimulus evokes spatiotemporal waves that percolate across the cortex. The spatial characteristics of this wave, however, are not readily apparent from just observing pairwise phase relationships. Thus, we examined the spatial characteristics of the visual evoked oscillations that are simultaneously recorded across multiple locations on the cortical surface. Trial average LFP filtered at fast and slow frequencies along the anterior-posterior (AP) axis recorded in a single representative mouse are shown in Figure 2a Sup. Video 3 and 2b respectively, (see Sup. Video 2, and Sup. Video 3 for propagation of visual evoked fast and slow waves, respectively, over the cortical surface). Oscillations observed at each electrode are consistently phase shifted in relation to oscillations at neighboring electrodes. Thus, the overall ensemble activity profile resembles traveling waves at both frequency bands. Remarkably, the fast wave is initiated in the visual cortex and propagates anteriorly, while the slow oscillation initiates rostral to V1 and spreads in the opposite direction (Figure 2C, Sup. Video 4). Note that travelling waves that propagate through uniform media have a uniform spatial phase gradient at all locations. Consequently, the phase offset ought to grow linearly with distance. This is approximately true of signals over short distances in Figure 2. In contrast, over long distances a clear nonlinear relationship between phase offset and distance is seen. This nonlinear relationship implies that the propagation of these wave-like patterns is likely to depend on the specifics of network architecture in different cortical regions.

**Figure 2:**
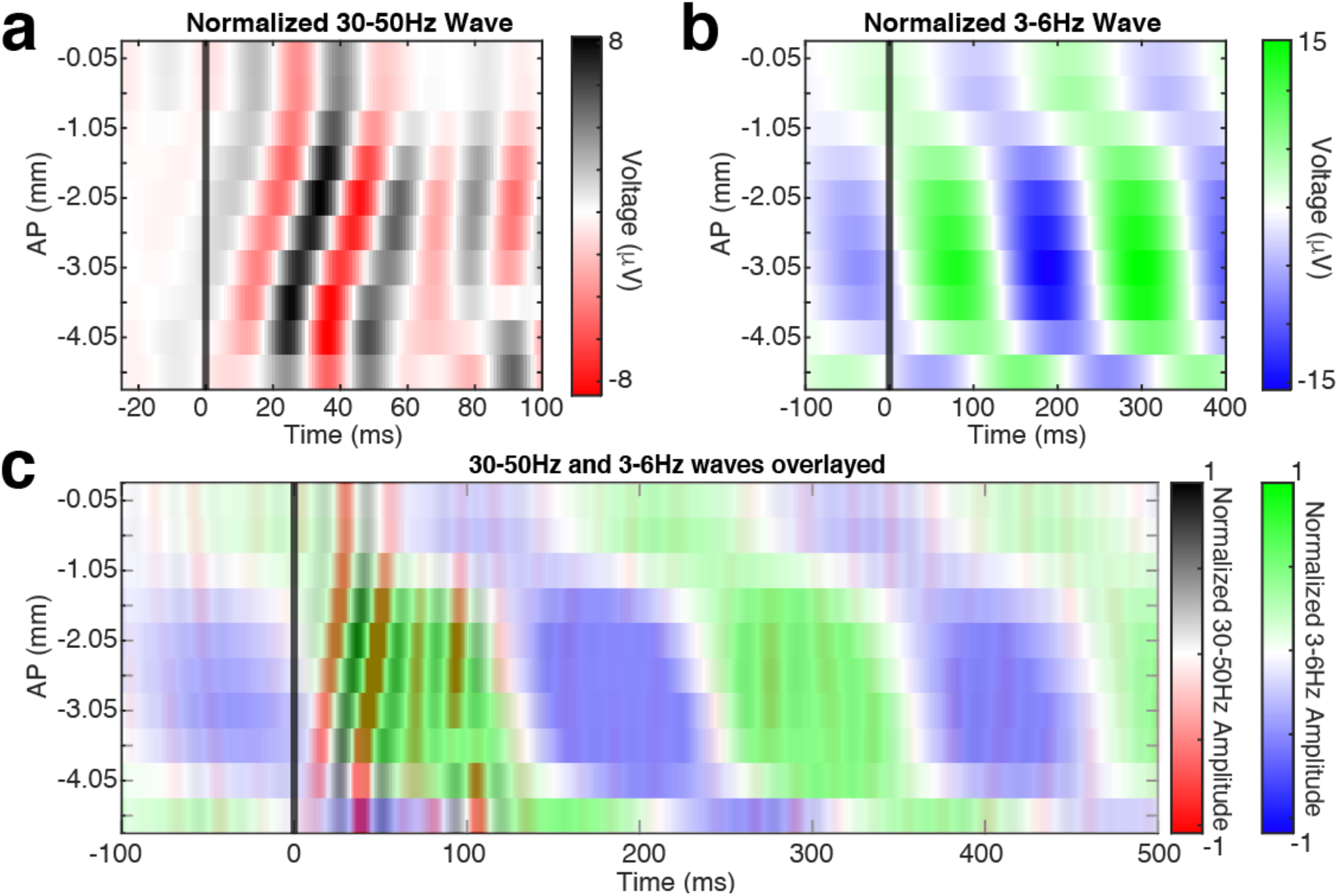
Average filtered LFP illustrate traveling wave-like behavior. a. Average of the VEP filtered at 30-50Hz from 10 electrodes along the anterior to posterior axis (−2.25mm ML) in a representative mouse. The x axis denotes time relative to stimulus onset, the y axis indicates AP position of an electrode relative to bregma. Note evoked high frequency waves starting at -4.05mm from bregma (V1) and traveling anteriorly over ∼100ms. b. Average of the VEP filtered at 3-6Hz from the same mouse arranged in the same format at B. Note the low frequency waves begin more anteriorly (∼2 mm from bregma) relative to the fast oscillations and travel in the posterior direction. c. Superimposition of the data in A-B (amplitude of the signals is normalized to highlight phase relationships between oscillations at different temporal frequencies). The fast wave begins posterior to the slow wave and travels rostrally towards the slow wave initiation zone.

### Coherent spatiotemporal waves are detected using complex SVD

While data in Figure 2 strongly suggest a propagating wave-like phenomenon, the interpretation of these data is somewhat limited. First, the LFP is a complex mixture of spontaneous and evoked activity (Arieli et al., 1996; Kisley and Gerstein, 1999). Second, trial averaging may obscure single trial behavior. Thus, to provide additional evidence that simple visual stimuli elicit travelling wave-like phenomena, we applied a methodology to separate spontaneous from evoked activity and to characterize spatiotemporal features of evoked activity on a single trial level. For this purpose, we utilized Singular value decomposition (SVD) of the complex-valued analytical signals derived from bandpass filtered LFPs.

Singular value decomposition (SVD) factorizes a spatiotemporal matrix into mutually orthogonal spatiotemporal modes:

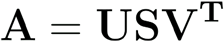

where ***A*** is an *n* by *t* matrix that contains *n* channels of analytical signals sampled at *t* time points, ***U*** is an *n* by *n* complex valued spatial matrix in which each column encodes the phase and amplitude of a single mode at each channel, ***V*** is a *t* by *t* complex valued temporal matrix in which each row encodes the instantaneous phase and amplitude of each mode at each time point. Finally, ***S*** is an *n* by *t* diagonal real valued matrix which encodes the fraction of the total signal contained in each mode.

The advantage of performing SVD on the complex-valued analytical signal is that projecting the data onto the complex plane linearizes phase relationships between channels. In contrast, phase-shifted real-valued oscillations across channels would exhibit correlations at different time lags and are therefore not easily factorizable using SVD or similar dimensionality reduction techniques. We highlight the utility of complex SVD with synthetic data in Sup. Figure 2.

Here, we performed SVD on the analytical signal of single trials filtered at fast and slow frequencies. 72% of variance of single trial VEPs was captured by the first ten singular modes (*95% Confidence Interval* = 62%-81%). We then defined the most visually responsive mode for each trial as the mode in which the post-stimulus temporal amplitude increases the most compared to pre-stimulus amplitude (Sup. Figure 3). The most visually responsive mode was most often associated with the largest singular value and thus contained the highest amount of signal variance. The results of the analysis are robust to the changes in the total number of modes considered for the analysis.

### Visual evoked waves have a consistent phase relationship from trial to trial and across animals

The spatial phases of the most visually responsive mode from each single trial were aggregated across trials and mice (Methods). The phase difference between visual evoked fast waves in two locations in V1 (black and red diamonds in Figure 3c and d) reveals a consistent phase offset across trials and mice (Figure 3a). As the distance from the V1 electrode is increased, the phase difference between the oscillations grows concomitantly (Figure 3b). Consistent with the average LFP data (Figure 2), a progressive increase in phase offset with distance suggests that the activity evoked by the visual stimulus on a single trial level has characteristics resembling a travelling wave. Furthermore, the tight phase offset distribution implies that the spatial properties of these evoked waves are highly consistent from trial to trial and between animals.

**Figure 3:**
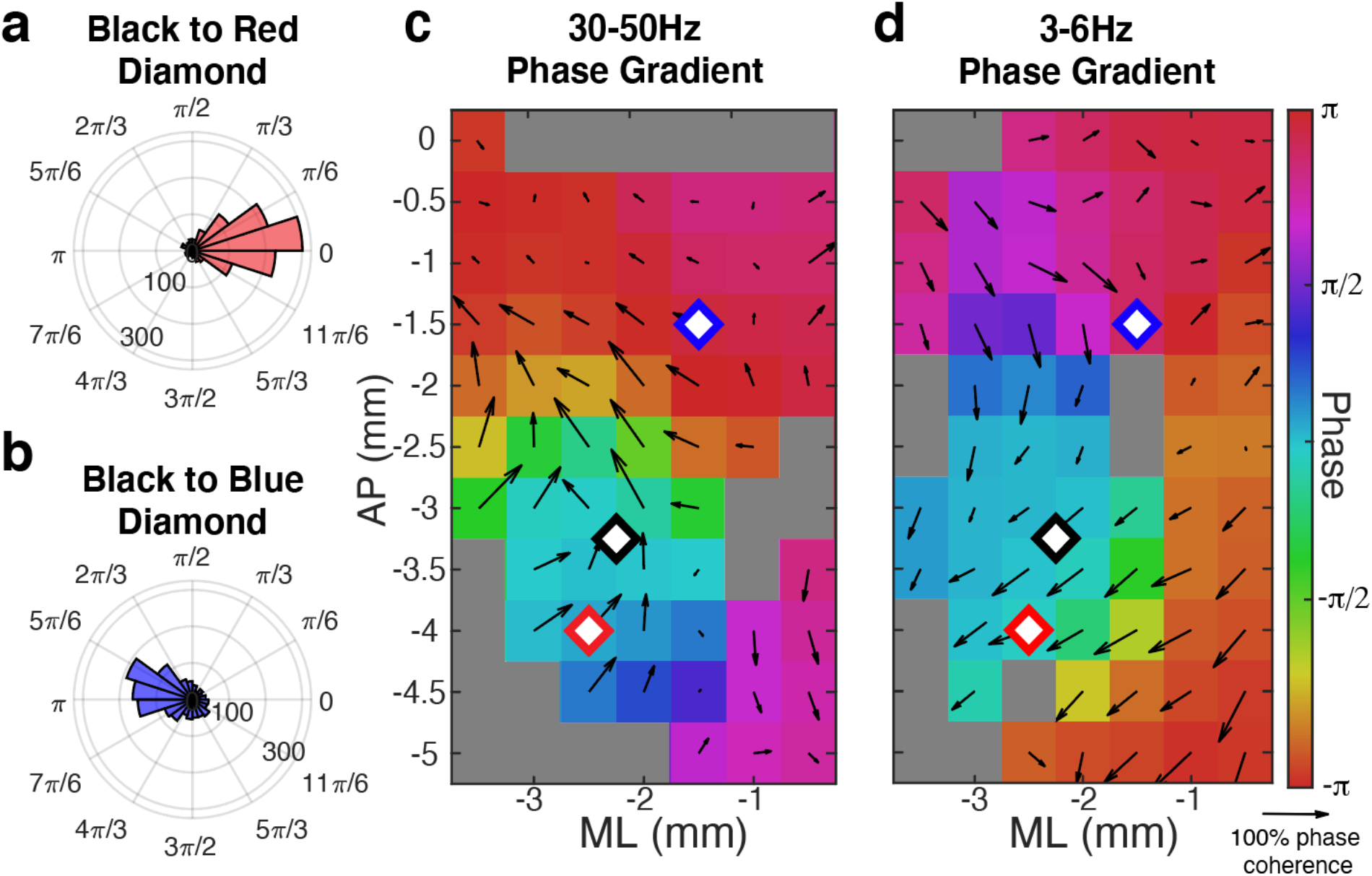
SVD identifies visual evoked traveling waves that are consistently elicited both across trials and across animals. a. Histogram of the difference in the spatial phase of the most visually responsive 30-50 Hz mode between two electrodes in V1 (black and red diamonds in C) across trials and animals. b. Histogram of phase angle difference of the most visually responsive 30-50 Hz mode between an electrode in V1 (black diamond in plot C) and PPA (blue diamond in plot C) across trials and animals. Note that the phase angle difference is increased as distance from V1 increases. c. At each stereotaxic location, the average phase offset of the 30-50Hz spatial mode relative to V1 (the black diamond) is plotted in color. Spatial phase gradient is depicted by black arrows. The direction of the arrows shows the direction of spatial phase gradient over trials and mice. The length of the arrows is 1-circular variance and therefore corresponds to the consistency of the angle of the spatial phase gradient over trials and animals (scale arrow for 100% phase coherence underneath color axis in D). Locations that are greyed out did not meet Bonferroni corrected statistical significance (p-value < 0.0006, Raleigh test). d. Phase offset relative to V1 and the spatial phase gradient of the most visually responsive 3-6Hz spatial mode at each stereotaxic location depicted as in D.

The spatial phase of the most visually responsive mode averaged across trials and animals is shown in Figure 3c and 3d for the fast and slow oscillations, respectively. This confirmed that throughout most of the cortical surface, the phase relationship between evoked fast and slow oscillations is consistently observed from trial to trial and among animals. Consistent with the example observed in the average filtered signal (Figure 2), the phase gradient for fast and slow oscillations evolves in approximately opposite directions. Thus, a brief visual stimulus elicits both fast and slow spatiotemporal activity patterns that resemble travelling waves and percolate over the cortical surface for hundreds of milliseconds. The fast wave propagates in the feedforward direction from the visual cortex towards higher order cortical areas. The slow wave propagates in the feedback direction from the higher order cortices back towards the primary visual cortex. Given the initiation zones and directions of propagation, we will refer to the fast visual evoked wave as “feedforward” and the slow visual evoked wave as “feedback.”

Similar feedforward and feedback propagating waves were observed for weaker visual stimuli (Methods). For weaker stimuli, the propagation of the fast visual evoked waves was predominantly limited to V1 and was not affected by the stimulus intensity. In contrast, the spatial extent of the feedback slow visual-evoked wave strongly depended on stimulus intensity. For lowest luminance stimulus, the feedback slow wave was principally observed in V1. However, for higher luminance stimuli, the feedback travelling slow wave involved much of the cortex (Sup. Figure 4). Thus, the spatial extent of the feedback wave tracks stimulus intensity, in a manner that mirrors psychophysics (Wysocki and Stiles, 1982; Denman et al., 2018).

### High and low frequency waves are present throughout the cortical layers in V1 but are constrained to the superficial layers of the posterior parietal cortex

To identify the circuits mediating the visual evoked waves, we computed current source density (CSD) from the two laminar probes targeting V1 and PPA (Figure 4a, 4b, 4f). In V1, we identified a canonical CSD pattern. The first sink occurs in the granular layer. Subsequently, alternating sink and source patterns occur throughout the cortical column, revealing communication among the cortical layers (Figure 4c). Less is known about the neurophysiological responses of the PPA to visual stimuli. We find the first sink at 0.15 mm below the cortical surface, which appears at a longer latency than in V1. Moreover, the majority of the CSD signal in PPA is confined to the superficial layers (Figure 4g).

**Figure 4:**
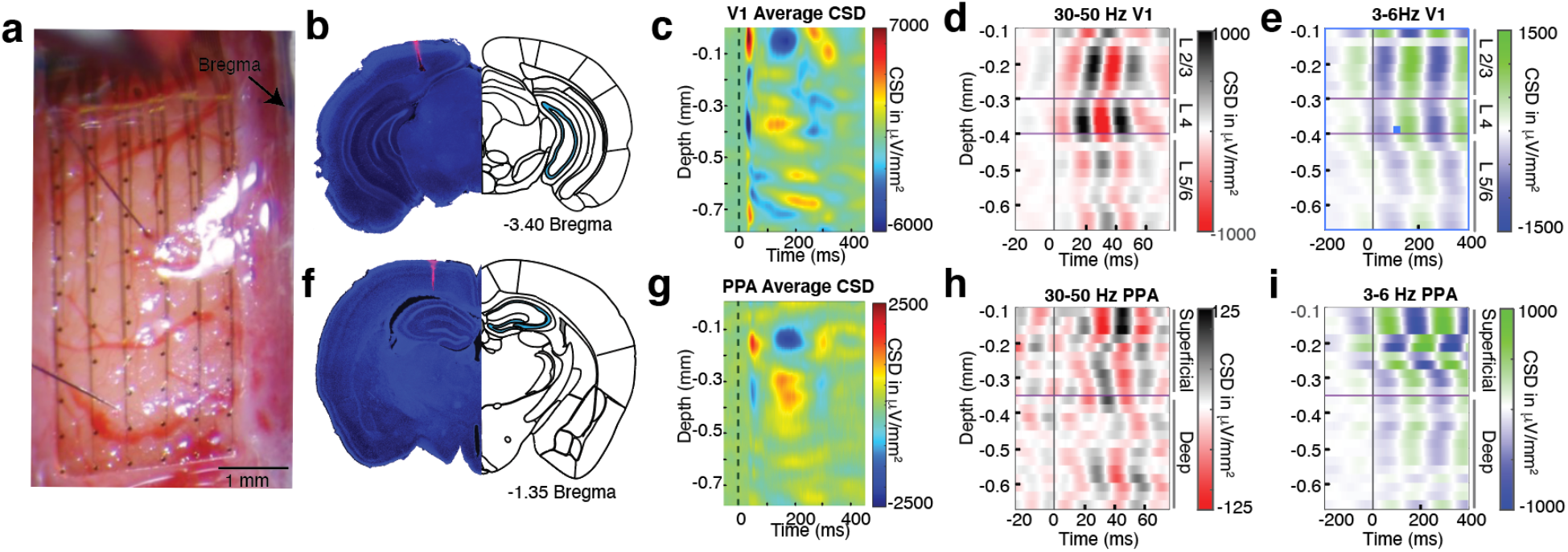
Intra-laminar recordings reveal. a. Photograph of 64 channel electrocorticography grid with two 32 channel penetrating laminar probes placed in through holes in the grid into V1 and PPA. b. Histological verification of laminar electrode localization in V1. c. Current source density (CSD) in V1 averaged over trials and mice. d. V1 CSD filtered at 30-50Hz and averaged across trials in a representative mouse. 30-50Hz oscillations originate in layer IV and then propagate to superficial and deep cortical layers. In D and E purple lines show approximate location of layer IV defined by the earliest sink in the CSD. e. Same as D but filtered at 3-6Hz. The 3-6Hz oscillations appear approximately simultaneously in the supra- and granular layers and propagate to deeper layers. f. Histological verification of laminar electrode localization in PPA. g. CSD in PPA averaged over trials and mice. Purple lines show approximate location of superficial layers I-IV, and deep layers V/VI based on anatomy (Allen Brain Atlas). h. Same as D for PPA. The 30-50Hz oscillations are most prominent in the superficial layers. i. Same as E for PPA. The 3-6Hz oscillations begin in the superficial layers and propagate to deeper layers *Data in D, E, H, I are from the same representative mouse.

Frequency domain analysis reveals strong ITPC for the fast frequency at all cortical layers in the first 100 ms following the stimulus in V1. A similar pattern is observed for the slow oscillation for ∼ 500 ms after the stimulus (Sup. Figure 5a, 5c). Within the PPA, in contrast, most of the ITPC at both high and low frequencies is concentrated in the superficial cortical layers (Sup. Figure 5b, 5d).

To determine the laminar organization of fast and slow waves, we averaged the filtered CSD data at each depth within each mouse. Consistent with other work on visual evoked gamma oscillations in V1, the fast waves originate in layer 4 in V1 and propagate to supra- and infra-granular layers (Figure 4d), indicating a critical role of thalamocortical circuitry in the initiation of the visual evoked gamma oscillations (Reinhold et al., 2015; Saleem et al., 2017). In contrast, in the PPA, visual evoked fast oscillations are predominantly seen in the superficial layers (Figure 4h). The visual evoked slow oscillations originate in the superficial layers in both V1 and PPA (Figure 4e, 4i). These observations imply that the fast visual evoked waves are initiated through the interactions between the thalamus and the input layer 4 of V1 and subsequently propagate through the corticocortical circuitry involving supra- and infra-granular layers in the feedforward direction towards higher order cortices (Feldmeyer et al., 2002, 2006; Douglas and Martin, 2004, 2007; Krieger et al., 2007; Wester and Contreras, 2012). In contrast, the slow visual evoked wave predominantly propagates in the ventral direction through the cortical column, supporting the conclusion that it is primarily mediated by the feedback cortico-cortical interactions.

### Both fast and slow visual evoked waves have large spatial frequencies

It is commonly thought that waves with higher frequency tend to be localized in space, whereas slow temporal frequency waves involve large areas of the cortex (Kopell et al., 2000). In contrast to these observations, we show that both the fast and the slow oscillations involve much of the cerebral hemisphere for supra-threshold stimuli. Further, examination of the recordings in Figure 2a and 2b suggests that despite their difference in temporal frequency, the spatial wavelengths of both waves are similar. We confirm this observation and estimate the most common spatial wavelengths of the fast and slow waves to be 12.7 mm/cycle and 12.5 mm/cycle, respectively (Figure 5a). These spatial wavelengths are on or above the scale of a mouse cerebral hemisphere (Ermentrout and Kleinfeld, 2001). Because of the nonlinear dependence of phase offset on distance (Figure 2) each travelling wave does not have a well-defined single spatial wavelength. Nevertheless, local estimates of spatial wavelengths can be obtained from the spatial phase gradient (Figure 3c, 3d) at each cortical site (Sup. Figure 6a, 6b). Thus, while visual evoked fast and slow waves are distinct from canonical travelling waves in uniform medium and do not have a single spatial wavelength, the spectra of spatial wavelengths for the fast and slow oscillations are comparable. The propagation velocity of the fast oscillations, consequently, is approximately an order of magnitude faster than the slow oscillation (median_fast_= 0.8 m/s, IQR_fast_= 0.5-1.58 m/s, and median_slow_= 0.11 m/s, IQR_slow_= 0.07-0.20 m/s, for the slow and the fast waves respectively). This, again is consistent with data in Figure 2. The differences in the propagation velocities suggest that the fast and slow visual evoked waves are mediated by different circuit mechanisms.

**Figure 5:**
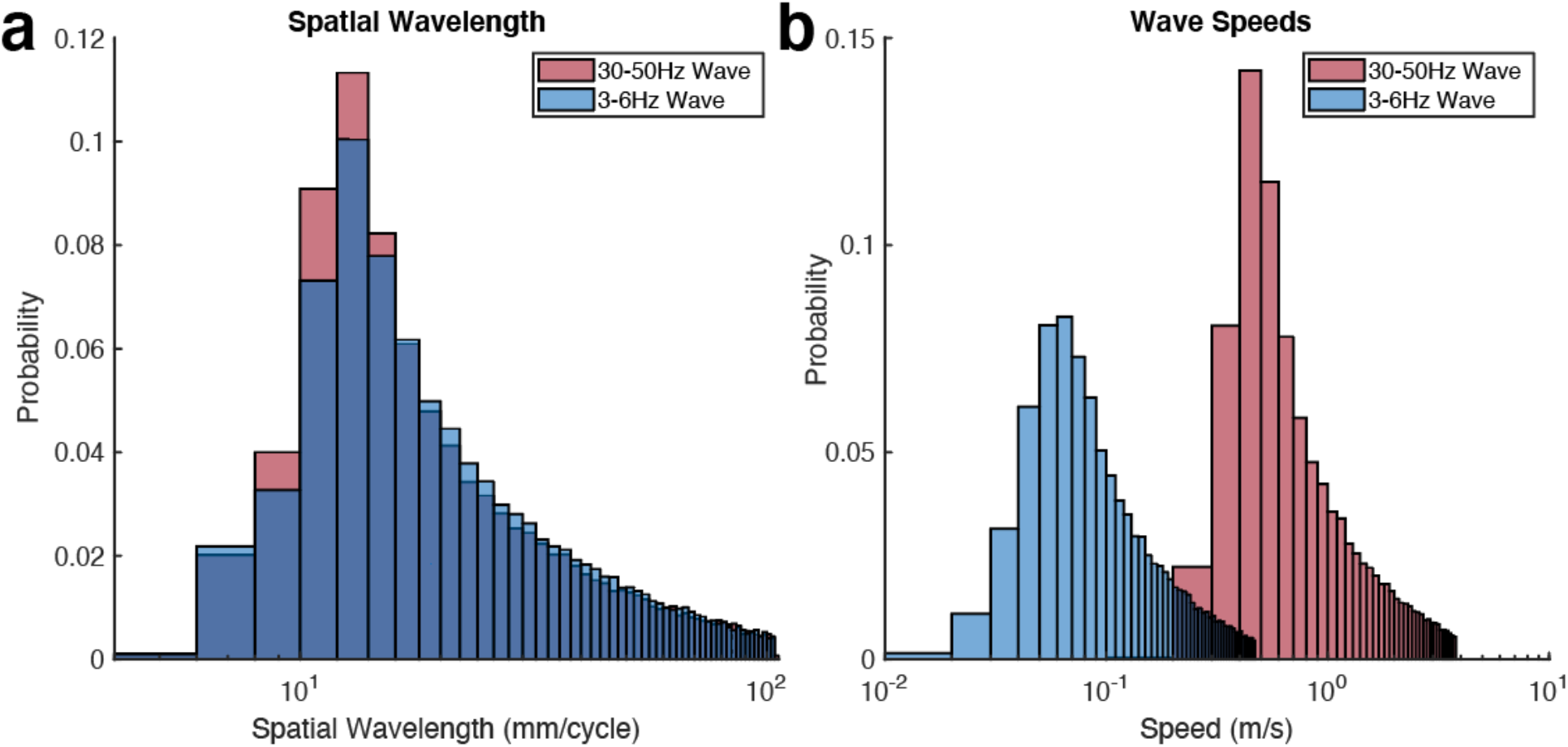
Fast and slow visual evoked waves have similar spatial wavelengths but significantly different propagation velocities. a. Distribution of spatial wavelengths of 30-50Hz and 3-6Hz most visually responsive modes. b. Distribution of wave speeds of most visually responsive modes at 30-50Hz and 3-6Hz.

### Fast and slow oscillations comprise a single multiplexed visual evoked spatiotemporal response

Until this point, we have treated the high and low frequency visual evoked waves as independent entities. Furthermore, we only considered the waves observed in the immediate aftermath of the stimulus. However, analysis of single trials in V1 reveals rhythmic waxing and waning of the amplitude of fast oscillations aligned to the phase of the slow oscillation (Figure 6a). Similar phase amplitude coupling is also observed in the PPA (Figure 6b); although the amplitude of fast oscillations peaks at different phases of the slow oscillation. Indeed, significant phase amplitude modulation is present throughout the cortical surface (Figure 6c). Moreover, the phase relationship varies systematically with cortical location (Figure 6d). Thus, the fast and slow waves are not independent phenomena, but instead are different aspects of the same integrated spatiotemporal activity pattern, which is reliably evoked by the visual stimulus.

**Figure 6:**
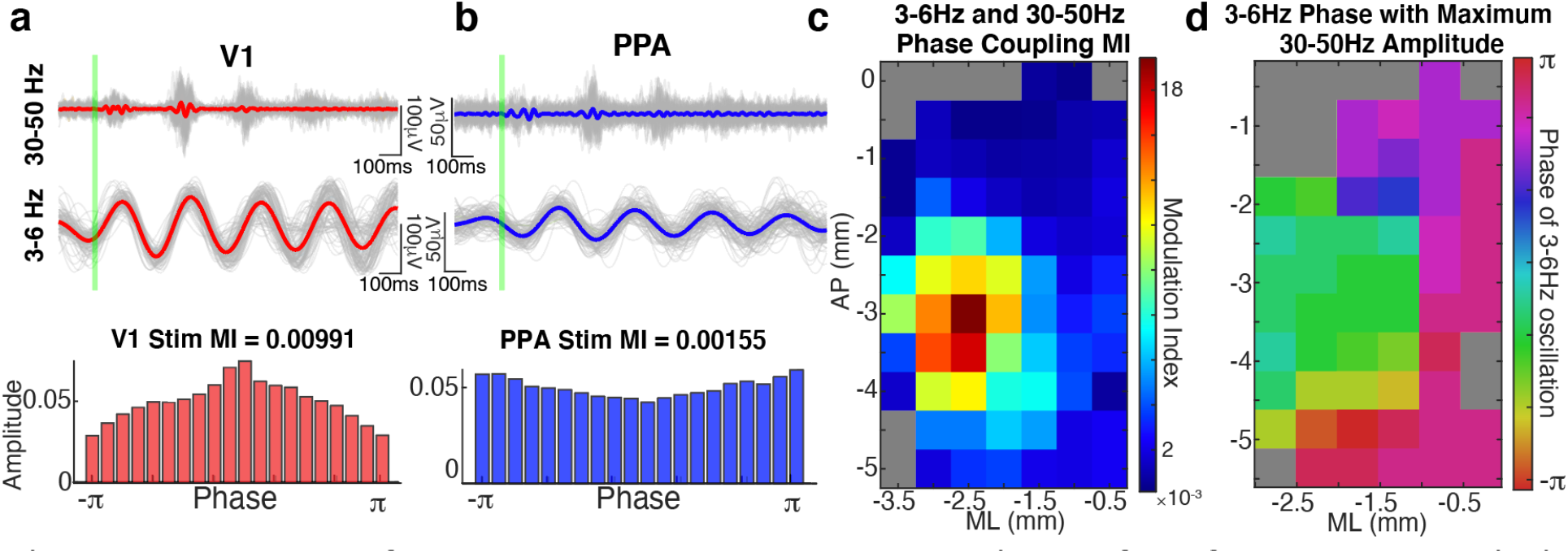
The phase of the slow wave modulates the amplitude of the fast wave both within V1 and throughout the cortex. a. Top: single trials (gray) and average (red) data filtered at 30-50Hz over V1 of a representative mouse. Middle: same as above, but for 3-6Hz. Note fast oscillation bursts occur rhythmically in phase with slow oscillations. Bottom: Amplitude of high frequency oscillations at each phase of the low frequency oscillation, averaged over trials. The deviation of this distribution from a uniform distribution is summarized in the modulation index (MI) of 0.00991(p-value <0.0001, compared to time shuffled surrogates). b. Similar to A but for an electrode over PPA. The phase amplitude MI = 0.00155 (p-value <0.0001). Note that the phase of the 3-6Hz oscillation at which the gamma amplitude is maximum is shifted compared to that in V1. c. Modulation indices averaged over all 13 mice and plotted in color at each stereotaxic location. Locations that are greyed out did not meet Bonferroni corrected statistical significance (p-value < 0.0006) compared to time shifted surrogate data. MI peaks near V1 but remains statistically significant over much of the cortical surface. d. The phase of the slow 3-6Hz oscillation at which the fast 30-50Hz oscillation reaches maximum amplitude as shown for a representative mouse at each stereotaxic location. Grayed out locations did not meet statistical significance compared to time shifted surrogates.

### The phase of slow visual evoked waves modulates the firing rates of neurons both in V1 and PPA

Fast oscillations in the gamma range are thought to coincide with neuronal firing. In contrast slower oscillations are dominated by synaptic potentials (Buzsáki et al., 2012). The phase amplitude coupling between the fast and slow visual evoked oscillations may therefore suggest that the slow feedback oscillation modulates neuronal firing. To determine whether this is indeed the case, we tested whether the slow visual evoked waves entrain firing of single units in V1 and PPA. We first isolated single units throughout the cortical lamina in V1 and PPA (155 in PPA and 186 in V1, Figure 7a and 7b for representative neurons in each area, respectively). Raster plots of these neurons (Figure 7c and 7d) show that after the stimulus the firing of the neurons in both areas is entrained by the slow visual evoked oscillation. To quantify this observation, we computed spike field coherence for each single unit and the CSD filtered at the slow frequency band from the same lamina. 32 out of 155 units in PPA and 98 out of 186 units in V1 exhibited significant spike field coherence after the stimulus (Figure 7e and 7f). Spike field coherence for the same units was significantly smaller before the stimulus (*p<10*^*-5*^ *Mann-Whitney U-test)*. Thus, visual evoked slow oscillations entrain a significant fraction of neurons both in the primary visual cortex and the association cortex. While many single units were entrained in both cortical areas, the phase of maximum firing was not the same across different units (Figure 7e and 7f). Indeed, the phase of maximum firing in each area swept through an entire cycle of the slow wave. Thus, each visual stimulus evokes a sequence of neuronal activation in both areas that is orchestrated by the slow oscillation.

**Figure 7:**
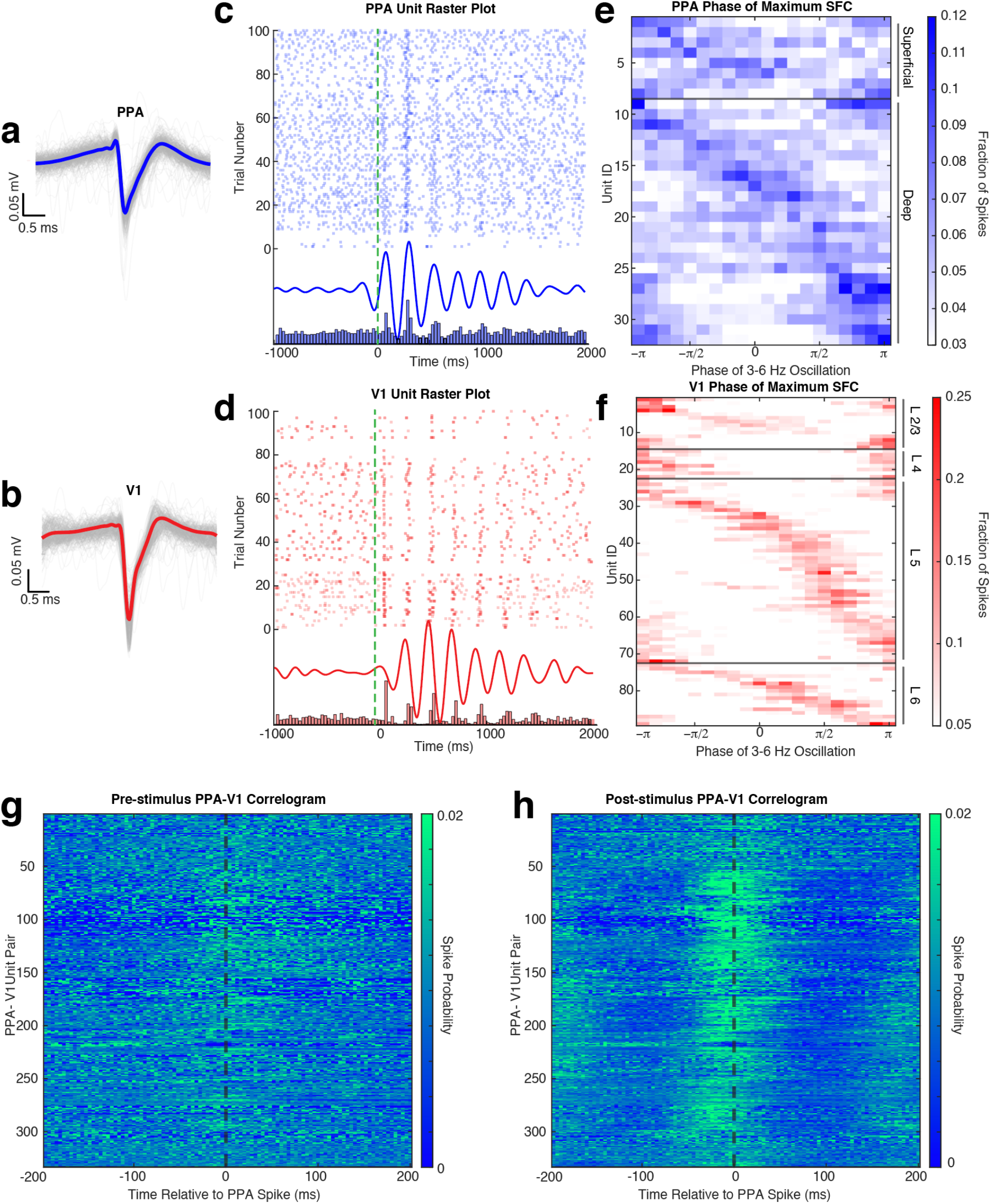
Probability of neural spiking in both V1 and PPA depend on the phase of the slow wave. a. Individual action potential waveforms of a representative PPA unit located in layer V (gray traces), with the average waveform superimposed in blue. b. Same as A, but for a representative V1 neuron located in infragranular layers (gray traces), with the average waveform superimposed in red. c. Raster plot (top) of 100 trials of PPA unit (green dashed line marks stimulus onset). The average CSD, filtered at 3-6 Hz, of the LFP at the same depth as the unit in A (middle). The peristimulus histogram of the same unit (bottom). d. Same as C, but for the representative V1 unit in B. e. Probability of firing as a function of phase of the slow oscillation for each unit in the PPA. Each row is an individual unit in the PPA that has statistically significant spike field coherence (SFC) with the slow oscillation. Units above the black horizontal line are in the superficial layers of the PPA. Units below the black horizontal line are within the deep layers. f. Same as E for V1 units with statistically significant spike field coherence. The horizontal back lines highlight four sections in which V1 cells reside, in top-down order: layer II/II, layer IV, layer V, and layer VI. g. Cross-correlograms between PPA and V1 neurons entrained by the slow wave during the 500 ms before visual stimulation. Each row is an individual PPA V1 pair. Probability of firing is shown by color. h. Same as in G but for 500 ms after the stimulus.

If the visual evoked waves were responsible for coordinating neuronal activity across disparate regions in the cortical hierarchy, one would expect that neuronal firing would become transiently correlated after stimulus presentation. Consequently, we hypothesized that V1 and PPA neurons that are entrained by the slow wave would become transiently correlated after the stimulus. As expected, prior to the stimulus, firing in V1 and PPA was largely uncorrelated (Figure 7g). However, after the stimulus, many of these previously independent neurons became correlated over half of the wave cycle length of the slow wave (∼100ms) (Figure 7h). Thus, as the feedback slow visual evoked wave propagates from the higher order cortical areas towards the primary sensory cortex, it entrains a sequence of neuronal activation in the PPA and V1. This provides a neurophysiological insight into how simple sensory stimuli produce coordinated patterns of neuronal activity that span multiple cortical areas.

## Discussion

Here, we show that in awake mice, a brief presentation of a simple visual stimulus reliably evokes two traveling waves. The spatiotemporal characteristics of these waves are highly stereotyped across individual trials and across animals. Fast (30-50Hz) waves begin in layer 4 of V1 and travel anteriorly in a feedforward manner. Slow (3-6Hz) waves are initiated in the superficial layers of the higher order areas and travel posteriorly in a feedback fashion. These waves are tightly coupled forming a single multiplexed spatiotemporal wave-like activity pattern observed throughout the cortex. The phase of the feedback wave modulates the firing of individual neurons both in the association cortex and in V1. A consequence of this entrainment is that previously independent neurons in V1 and PPA form a transient coordinated assembly after stimulus presentation. In this way, the feedback and the feedforward aspects of the multiplexed visual evoked waves coordinate neuronal activity across distant cortical regions involved in the processing of visual stimuli.

The role of neuronal oscillations in mediating feedforward and feedback sensory processing has been predominantly studied by analyzing pairwise signal covariation. Our chief contribution is that a set of such pairwise coupled neuronal oscillations together form a single coherent spatiotemporal pattern that consists of two interacting waves. Spontaneous and stimulus evoked travelling waves have been observed in the EEG (Adrian and Matthews, 1934; Goldman et al., 1948; Adrian and Yamagiwa, 1960; Darrow and Hicks, 1965; Hughes, 1995). However, the interpretation of the EEG is hindered by low spatial resolution and volume conduction. Further, intracranial recordings (ECoG) did not corroborate the EEG findings (Hangya et al., 2011; Bahramisharif et al., 2013; Mak-McCully et al., 2015; Muller et al., 2018), leaving the existence and functional role of macroscopic traveling waves in question. Novel experimental imaging techniques using voltage sensitive dyes (VSDs) reveal mesoscopic travelling waves that are confined by anatomical boundaries between cortical regions (Polack and Contreras, 2012; Muller et al., 2014) or produce complex interference patterns at the inter-region boundaries (Xu et al., 2007b). However, most mesoscopic waves recorded with VSDs do not take into consideration the temporal frequency of travelling waves and focus primarily on their spatial propagation properties (Benucci et al., 2007; Xu et al., 2007a; Polack and Contreras, 2012; Sato et al., 2012; Muller et al., 2014, 2018; Townsend et al., 2015; Davis et al., 2020). Here, by identifying two temporal frequencies that are reliably phase locked to the stimulus, we deconstruct the overall spatiotemporal response pattern into two distinct travelling waves that percolate through the brain in opposite directions. This, in turn, allows us to experimentally marry pairwise feedforward-feedback interactions involving different temporal frequencies and travelling cortical waves into a single, unified framework.

Feedforward and feedback aspects of sensory processing serve fundamentally different roles. Feedforward processing assembles increasingly abstract representations of sensory stimuli. Feedback processing, in contrast, situates sensory stimuli within a behavioral context. Based on these functional differences, one expects that the neurophysiological processes that mediate feedback signaling must occur on a slower time scale than those involved in feedforward interactions. This assertion is consistent with experimental work in primates (Engel et al., 2001; Bosman et al., 2012; Womelsdorf et al., 2012; Van Kerkoerle et al., 2014; Bastos et al., 2015; Michalareas et al., 2016). Many studies demonstrate that the faster gamma oscillations underlie feedforward processing while the slower, alpha oscillations, relay feedback processing. This difference in time scales is confirmed by our experimental observations – the temporal frequency of the feedback wave is approximately ten times slower than the feedforward wave. While there is a numerical discrepancy between the temporal frequency of alpha oscillations in primates and the 3-6Hz feedback wave in our work, multiple lines of evidence strongly suggest that the 3-6Hz wave in mice is analogous to the primate alpha oscillations (Bollimunta et al., 2008, 2011; Dougherty et al., 2017; Einstein et al., 2017; Senzai et al., 2019; Speed et al., 2019; Nestvogel and Mccormick, 2021). Our identification of feedforward and feedback processes as interacting travelling waves permits an extension of this postulate. We find that the propagation velocity of the feedforward wave is also roughly an order of magnitude faster than that of the feedback wave. This difference in propagation velocities potentially allows the feedback modulation to integrate across recent sensory stimuli, thereby situating them in behavioral context.

We refer to the activity patterns evoked by visual stimuli as “traveling waves”. However, it is important to note that these large-scale spatiotemporal responses differ from simple waves in uniform medium. Imagine that a response to a visual stimulus is akin to a raindrop falling into a still pond. In this highly idealized case, the raindrop would create a wave radiating outward at uniform speed and spatial wavelength. This simple scenario is indeed similar to travelling waves within a single cortical area. Much like waves on the pond, travelling cortical waves typically have tight distributions of propagation speeds and spatial wavelength (Xu et al., 2007b; Muller et al., 2014; Besserve et al., 2015). Our results are in agreement with these findings over relatively short spatial scales (Sup. Figure 6). However, over larger scales, the apparent “viscosity” of the medium changes. There is a clear departure from the linear dependence of the spatial phase gradient on distance. This gives rise to a broad spectrum of spatial wavelengths and propagation velocities. The “viscosity” of the brain is thought to arise from conduction delays between different neuronal oscillators (Ermentrout and Kleinfeld, 2001). The observation that speed of wave propagation deviates from a pure travelling wave on large spatial scales in a systematic fashion suggests therefore that different conduction delays are involved on small and large scales.

It has been hypothesized that the interactions between distinct neuronal oscillators are mediated by horizontal fibers in superficial cortical layers. The spatial properties of traveling waves within a single cortical region are consistent with this hypothesis (Grinvald et al., 1994; Bringuier et al., 1999; Muller et al., 2014). The propagation speeds of the visual evoked travelling waves observed herein are also in the range of conduction delays of cortico-cortical fibers. Direct laminar recordings showing preferential involvement of superficial cortical layers provide additional evidence for this hypothesis. While on the scale of a single cortical region, wave propagation is likely predominantly mediated by horizontal cortico-cortical fibers, additional mechanisms likely contribute to propagation over large spatial scales. For instance cortico-thalamic and corticobulbar loops contribute significantly to processing of visual stimuli and involve superficial cortical layers (Burkhalter, 1989; Feldmeyer et al., 2002, 2006; Brumberg et al., 2003; Douglas and Martin, 2004, 2007; Wester and Contreras, 2012; Glickfeld et al., 2013; Townsend et al., 2017). The contribution of multiple anatomical pathways with distinct conduction velocities together with anisotropic connectivity likely distort the speed and direction of propagation of travelling waves on large spatial scales. This conjecture is supported by models of coupled oscillators which suggest that specific patterns of conduction delays strongly influence the spatial features of the wave-like phenomena (Jeong et al., 2002). These results suggest a refinement to the current mechanistic models of travelling wave phenomena. Investigation of the relationship between the underlying anatomy and propagation properties of visual evoked waves on macroscopic scale may enrich our understanding of the relationship between neural architecture and the coordination of neuronal activity across the hierarchy of the visual system.

Our results demonstrate that the visual evoked waves are attractive candidates for organizing the feedforward-feedback computations necessary for sensory processing. Nevertheless, the full contribution of evoked waves to the processing of visual stimuli is unclear for several reasons. First, while we note the relationship between stimulus intensity and extent of the feedback wave, we do not directly address the behavioral significance of visual evoked travelling waves. Second, it remains unknown whether evoked travelling waves contain information about specific features of sensory stimuli or the animal’s behavioral state. Our study specifically focused on simple unstructured stimuli to identify the dominant spatiotemporal patterns evoked by them. The waves that we have identified are likely mediated by volleys of synaptic potentials (Bringuier et al., 1999), which modulate excitability of individual neurons. It is likely that the specific features of the stimulus are encoded by specific subpopulation of neurons that are phase locked to these waves. Processing of visual stimuli is known to involve activity of neurons broadly distributed across the visual system. Thus, while future work should address the relationship between stimulus features and the specific neuronal populations modulated by the travelling wave, our results show how such distributed neuronal assemblies can be coordinated by a wave that percolates across the visual system.

Traditional theories of sensory processing treat cortical neurons as independent feature detectors. This theoretical framework has been tremendously successful in predicting responses of individual neurons to stereotypical visual stimuli presented in isolation. However, models that treat neurons as independent feature detectors account for just a small fraction of activity in naturalistic settings (Olshausen and Field, 2005), indicating that spatiotemporal interactions between neurons are critical for effective visual processing. Recently, features of traveling waves have been associated with prioritization of neuronal responses in the new eye position after a saccade (Zanos et al., 2015), and detecting weak sensory stimuli (Davis et al., 2020). Our results add to this burgeoning evidence by showing that simple unstructured stimuli elicit waves of activity that percolate across space and time in a highly stereotyped fashion and entrain firing of neurons in distant cortical regions. Thus, instead of treating individual neurons as quasi–independent feature detectors, new theories of sensory processing should consider patterns of neuronal activity arising during interactions with a natural world as a superposition of waves.

## Methods

### Animals

All experiments in this study were approved by Institutional Animal Care and Use Committee at the University of Pennsylvania and were conducted in accordance with the National Institutes of Health guidelines. All experiments were performed using 8 male and 6 female adult (12– 32 weeks old, 20–30 g) C57BL/6 mice (Jackson Laboratories). Mice were housed under a reverse 12:12 h, light: dark cycle, and were provided with food and water ad libitum. A total of 20 mice were used in this study. Inclusion criteria for mice included the following: 1) presence of visual-evoked potentials (as defined by the absolute value of the average LFP response exceeding 5 standard deviations of pre-stimulus data within 100 ms after stimulus presentation) 2) histological verification of depth recording sites. With this inclusion criteria, we present data from 13 mice.

### Headplate implantation and habituation

At least 2 weeks before recording, mice were chronically implanted with custom designed headpieces for head-fixation during awake recordings using standard methodology. Briefly, mice were anesthetized with 2.5% and maintained at 1.5% isoflurane in oxygen, and secured a stereotaxic frame (Narishige). Local anesthesia (0.25 ml of 0.625 mg bupivacaine) and antiseptic (Betadine) were applied. Periosteum was exposed and additional local anesthetic (0.25 ml of 2% Lidocaine gel) was applied. Bregma and lambda as well as the site of the future craniotomy (+1mm to -5 mm AP, +0.25 mm to +6 mm ML left of bregma) were marked. The exposed skull was scored and the headpiece was attached using dental cement (Metabond) and 3 skull screws. Cyanoacrylate adhesive (Loctite 495) was applied over any remaining exposed skull. Mice were given 0.5mg cefazolin and 0.125mg meloxicam, and 7 ml of normal saline SQ after surgery. Animals were left to recover for a week before starting the habituation protocol. Mice were habituated to head fixation with body restraint with visual stimuli gradually over the course of 4 days. By the end of day 4, mice tolerated awake head fixation and visual stimuli for 45 minutes uninterrupted without any apparent distress.

### Craniotomy

On the day of the experiment, animals were anesthetized with 2.5% isoflurane in oxygen, and maintained at 1.5% isoflurane with closed loop temperature control (37+/- 0.5 degrees C) for the remainder of the surgery. 0.625 mg bupivacaine was injected in the surrounding face and neck muscles in order to provide scalp anesthesia. Mice were also given 0.5mg cefazolin and 0.125mg meloxicam, 0.006 mg dexamethasone and 7 ml of normal saline SQ, before surgery. Craniotomy was drilled through the dental cement over the markings on the left hemisphere (+1mm to -5 mm AP, +0.25 mm to +6 mm ML of bregma). One of the securing screws on the right skull bone was chosen as the reference. A 64-electrode surface grid (E64-500-20-60, Neuronexus) was positioned over the dura (most medial and anterior electrode was positioned ∼ 1mm lateral and 1mm posterior to bregma). Two laminar 32 channel probes (H4, Cambridge Neurotech) were coated with DiI (Sigma-Aldrich) for post mortem histological localization. The probes were inserted through the hole in the ECoG grid closest to V1 (−3.25 AP, -2.25 ML) and PPA (−1.5 AP, -1.5 ML) using a motorized micromanipulator (NewScale Technologies). Electrodes were inserted 800 µm into the brain at a rate of 25 µm/min. V1 electrode position was verified with current source density analysis. The grid and exposed dura was then covered with gel foam soaked in mineral oil. Isoflurane was then turned off for at least 20 minutes. At the end of this period and prior to recordings, animals were whisking, moving limbs and blinking in a manner similar to habituation before recordings began, thus suggesting that they were awake. This was corroborated by online analysis of the ECoG. After visual stimulation and recording, animals were deeply anesthetized (5% isoflurane) and sacrificed. Brains were extracted and fixed in 4% paraformaldehyde (PFA) overnight prior to sectioning and histology.

### Histology

Brains was sectioned at 80*μ*m on a vibratome (Leica Microsystems). Sections were mounted with medium containing a DAPI counterstain (Vector Laboratories). Electrodes were localized using epifluorescence microscopy (Olympus BX41) at 4x magnification.

### Visual Stimulation

Visual stimuli consisted of a 10 ms flash of a green LED (650 cd/m^2^) separated by random intertrial intervals sampled from a uniform distribution between 3 and 4 seconds. The flash covered 100% of the mouse’s visual field. In a subset of animals, visual stimulation was performed using a CRT monitor (Dell M770, refresh rate 60Hz, maximum luminance 75 cd/m2, positioned 23 cm away from the mouse’s right eye, at an angle of 60% from the mouse’s nose, thereby covering 70% of the mouse’s right field of view) at varying luminance (2%, 11% 44%, 75%, 100% of maximum screen luminance). Flashes were 100ms long and presented in a random order at a random time interval between 3 and 5 seconds.

### Electrode registration

After identifying the histological location of the two depth probes in each mouse, and with prior knowledge of the ECoG grid dimensions (i.e., 6 columns, 11 electrode rows, electrode spacing of 500*μ*m, electrode diameter of 60*μ*m, hole diameter of 200*μ*m,), the position of each ECoG electrode was triangulated in the following fashion. The ECoG grid is a semiflexible plane. The location of the cortical probes in the electrode coordinate system was given by the through holes used for electrode insertion. The stereotaxic coordinates of the electrodes were established using post-mortem histology by comparison to the brain atlas (Paxinos). The cosine of the angle, *q*, between the laminar probe positions in the electrode and stereotaxic coordinates was computed. Each electrode on the ECoG grid was assigned a location based on the Euclidean distance from the two laminar probe sites. The resultant grid location matrix was then multiplied to a rotation matrix (R) to obtain the final electrode positions in stereotaxic coordinates.

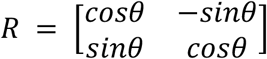

These coordinates were then verified by comparison to photographs of grid positions taken during experimental session.

### Electrophysiology and preprocessing

Signals were amplified and digitized on an Intan headstage (Intan, RHD2132) connected to an Omniplex acquisition system (Plexon, Omniplex), and streamed to disk at 40KHz/channel.

To extract LFP, data were downsampled to 1KHz and filtered offline using a custom-built FIR filter between 0.1Hz and 325Hz, with the MATLAB functions, *firls*.*m* and *filtfilt*.*m* to minimize phase distortion. Noise channels were manually removed and trials with excess motion artifact were rejected. Subsequently, the ECoG signals were mean re-referenced to minimize the effect of volume conduction. All further analysis was performed using custom built Matlab (Mathworks) code unless otherwise stated.

### Selection of electrode over Primary Visual Cortex (V1)

To average over animals, a single stereotaxic V1 location was selected as the electrode closest to (−3.25 AP, -2.25 ML), and in each animal. To confirm that the chosen electrode neurophysiologically corresponded to V1, the latency of onset of the VEP at each grid electrode was computed. The latency of onset of the visual-evoked potential was calculated as the time point at which if their post-stimulus average exceeds 3 standard deviations above the pre-stimulus baseline for 3 consecutive time points. The stereotaxically labeled V1 electrodes were 1-2 electrodes away from the electrodes with the earliest latency of onset in all mice and had latencies of onset within 2 ms of the electrodes with the earliest latency of onset.

### Current Source Density Analysis (CSD)

The one-dimensional CSD was computed as the second spatial derivative of the LFP recorded from the linear probes (Freeman and Nicholson, 1975):

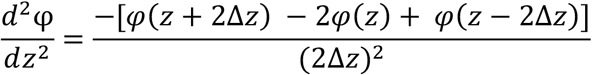

where ***φ*** is the LFP, z is the vertical coordinate depth of the probe, and Δz is the interelectrode distance (25 µm). CSD at the electrode boundaries were obtained using estimation procedure in (Vaknin et al., 1988). Cortical layers in the V1 probe were identified by the pattern of visual evoked current sinks and sources (colored as blue and red, respectively). Channels with the earliest current sink were assigned as layer 4 (granular layer). Subsequent sinks were found above and below layer 4 in layers 2/3 and layer 5. Laminar assignment of the channels in the PPA probe were based on distance from the cortical surface, where the CSD converged to zero. Channels within the first 350 µm were defined as superficial layers based on the thickness of layers 1-4 of the PPA (Reference Atlas :: Allen Brain Atlas: Mouse Brain; Franklin, Keith, B. and Paxinos, 2007; Lein et al., 2007). The next 400 µm were defined as deep layers. All further analysis of laminar LFP data was performed on the CSDs. V1 probe data was included only if there was a clear layer 4 sink and subsequent layer 2/3 and layer 5 sinks. Similarly, PPA probe data was included in analysis only if the superficial loss of CSD was seen, indicating that the most superficial electrode was positioned at the cortical surface. 11 mice fulfilled these criteria and were included in the analysis of laminar recordings.

### Spike Sorting

Single unit identification was performed on probe data from the same 11 mice that fulfilled criteria for laminar analysis. Spike sorting was performed using Kilosort (Pachitariu et al., 2016). The resulting spikes were then manually inspected for correct waveform clustering using Phy. All units with a firing rate lower than 0.5 spikes/s were excluded from further analysis.

### Wavelet analysis

Power, phase, and frequency information was extracted using a continuous wavelet transform using Morlet wavelets (0.1Hz to 150Hz, with a step-width 0.25Hz and normalized amplitude) (available at: http://paos.colorado.edu/research/wavelets/) (Torrence and Compo, 1998).

### Inter-trial Phase Coherence (ITPC) Analysis

Inter-trial phase coherence (ITPC) was used to quantify the phase synchrony between trials at each point in the time-frequency plot. ITPC was calculated for each electrode in each mouse. Briefly, angle vectors were extracted from the wavelet coefficients at each time point and each frequency by applying Euler’s formula and setting the single trial vector length to 1. ITPC was then calculated by taking the mean length of the angle vector across trials (Cohen, Mike, 2014).

ITPC values over the grid in Figure 1 were computed by averaging the ITPC over the first 100ms of the signal and over the frequency bands between 30-50Hz, or over the first 800ms of the signal and over the frequency bands between 3-6Hz

### Filtering data

LFP or CSD data was filtered into high (30-50Hz) or low (3-6Hz) frequency bands in order to perform phase based analysis. Data was filtered using the inverse wavelet transform, *invcwt*.*m*, (available at: http://paos.colorado.edu/research/wavelets/) (Torrence and Compo, 1998), by setting all wavelet coefficients outside the desired frequency range to zero.

### Analytical Signal Extraction

Hilbert transform was used to derive the analytical signal of LFPs or CSDs filtered in the gamma (30-50Hz) and low frequency (3-6Hz) data. This produced a time series of complex numbers. The modulus of the analytical signal is the instantaneous amplitude while the instantaneous phase is given by its arctan.

### Complex Singular Value Decomposition (SVD)

LFP recorded during a single trial and filtered at the appropriate frequency range (see above) using the Hilbert transform to derive an *n* × *t* analytical signal matrix ***A***, where *n* is the number of electrodes and *t* is the number of time points. Spatiotemporal modes were extracted from ***A*** by performing singular value decomposition which factorizes *A* into mutually orthogonal modes:

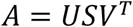

The columns of complex-valued *U* and *V* are the left and right singular vectors, which encode the spatial and temporal components of each mode, respectively. The diagonal real-valued *S* contains singular values (λ′*s*). The fraction of the total signal explained by *i* − *th* mode is given by

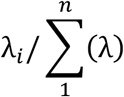

The spatial distribution of each mode is computed as *w*_*s*_ = |*U*_(∗,*i*)_| ∗ λ_*i*_. Each of the *n* components of *w*_*s*_ reflects the contribution of each electrode to the mode. The spatial phase is defined by θ_*s*_ = arctan *U*_(∗,*i*)_. Similarly, each component of θ_*s*_ reflects the spatial phase of each electrode. Temporal phase θ_*t*_ and amplitude *w*_*t*_ are defined in a similar fashion from **V**, θ_*t*_ = arctan *V*_(*i*,∗)_ and *w*_*t*_ = |***V***_(***i***,∗)_ | ∗ *λ*_*i*_. The spatial phase gradient of each mode 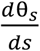 can then be computed locally at each electrode as in (Muller et al., 2014) (see below). The temporal frequency is given by the time derivative of the unwrapped temporal phase 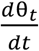 normalized by 2π. The analytical signal corresponding to mode *i* can be reconstructed as 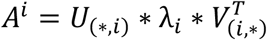. Finally, the LFP signals corresponding to each mode can be reconstructed as *LFP*^*i*^ = |*A*^*i*^| ∗ *cos*(arctan(*A*^*i*^)) . Illustration of this procedure is shown in Supl. Figure 2.

### Defining the most visually responsive mode

The first ten singular modes (accounting for 62.34 % - 81.18% variance, 95% confidence interval) computed for each single trial, as above. *w*_*t*_ is defined as the temporal amplitude for the first ten modes. *w*_*t*_ during the pre-stimulus period (400 ms) and was then used to compute the mean, ⟨w_*t*_⟩ and the standard deviation, 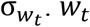 for the entire trial period (pre-and post-stimulus) was expressed as a z-score 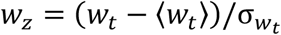. The most visually responsive mode was defined as the mode that exhibited the greatest increase in amplitude during the post-stimulus period (defined as 350ms post-stimulus for fast oscillations and 1000ms post-stimulus for slow oscillations).

### Spatial phase offset from V1

To determine the consistency in the phase relationship between spatial modes identified in different trials and across animals, the average difference in phase from each channel to the V1 channel was computed for the most visually responsive spatial mode. The V1 channel in each animal is defined as the channel closest to (−3.25 AP, -2.25 ML). The phase offset from V1 is calculated at each electrode by extracting the spatial phase of the most visually responsive mode θ_*s*_ and setting the V1 phase to zero. Circular mean and variance of *θ*_*s*_ referenced to V1 was then computed across trials and across animals (Fisher, N, 1995). The direction of the resultant vector corresponded to the average phase, whereas the magnitude of the vector is 1-circular variance.

### Spatial phase gradient

The spatial phase gradient was quantified by multiplying the complex spatial component of the most sensory evoked mode of a single trial at each electrode location to the complex conjugate of the spatial loading of its adjacent electrode, using continuous wavelet transform, *phase_gradient_complex_multiplication*.*m*, in MATLAB, written by Lyle Muller (available at: https://github.com/mullerlab/wave-matlab)(Muller et al., 2014). This operation was performed iteratively along the AP and ML direction of the grid. The resulting vectors were then converted into polar coordinates. To quantify the average gradient over trials, each trial’s gradient vector at each location was projected on a unit circle and the angles were averaged as above. Now, the angle of the resultant average vector is the direction of the average phase progression, whereas magnitude is 1-variance of the gradient over trials.

### Spatial Wavelength

The spatial frequency was computed by multiplying the magnitude of the single trial spatial gradient vectors and dividing the result by 2*π* to convert the units into cycles/mm. Spatial wavelength was calculated as the reciprocal of spatial frequency.

### Velocity of visual evoked waves

The instantaneous temporal frequency was computed by measuring the slope of the unwrapped temporal phase of the most visually responsive mode. The instantaneous frequency was then divided by the magnitude of the phase gradient of the most responsive spatial mode on a single trial basis.

### Phase amplitude coupling

Phase-amplitude coupling between oscillations was assessed using the modulation index measure (MI) of single trial filtered LFP data at every grid electrode site *REF to Tort. Phase of the 3-6Hz filtered data and the amplitude of the 30-50Hz filtered data were extracted from the analytical signal *A* as described above. Phase was binned into 20 phase intervals. The mean 30-50Hz amplitude was calculated for each bin for the first 900 ms of post stimulus activity per trial. The mean amplitude per bin was then averaged over trials. The MI was calculated by measuring the divergence of the resulting amplitude distribution from the uniform distribution using a modified Kullback–Leibler (KL) distance metric, with the function, *ModIndex_v2*.*m*, (available at https://github.com/tortlab/phase-amplitude-coupling) (Tort et al., 2010).

### Spike Field Coherence (SFC)

The phase 3-6 Hz filtered CSD was extracted from the analytical signal and segmented into 20 bins. The mean spike count was calculated for each bin for the first 900 ms of post stimulus activity per trial, and then averaged over trials. The MI was calculated by measuring the MI in a similar manner to calculating phase amplitude coupling.

### Averaging signals over stereotaxic coordinates

A query grid of stereotaxic locations was defined spanning -3.5 to 0 mm ML and -5 to 0 mm AP with 0.5mm spacing. For each query location, the weight of each electrode was assumed to depend on the distance between the electrode and the query location. The weights were defined to be a Gaussian function of Euclidian distance from the query location as follows:

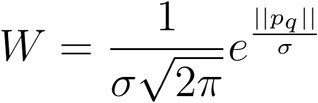

where │| *p*_*q*_ |│ is the Euclidean distance between the electrode p and query location q, σ = 0.15 is the standard deviation. The weight was computed as in the above equation for all electrodes within 0.3 mm of the query location and was set to zero otherwise. This weighting was used when computing averages and variances across mice as a function of stereotaxic coordinates.

### Spike Correlations

For each entrained V1 and PPA neuron in each animal, the delay of the spike times of the V1 cells relative to the spike times of PPA cells was computed during the pre-stimulus timeframe (500 ms before the stimulus onset), and post-stimulus time frame (0-500ms after stimulus onset). The distribution of spike time delays were then displayed in Figure 8.

### Statistical Analysis

To establish statistically significance of ITPC and phase coupling at each query location, the observed ITPC, MI values, and spike field coherence values were compared to a series of random time shifted surrogates (n = 100 sets, 100 trials per set). To determine the aggregate p-value over mice at each stereotaxic location, a Stouffer’s Z-score was calculated.

To determine if the phase relationships between each stereotaxic location and V1 and spatial gradient show a statistically significant deviation from a uniform distribution on a circle, a Rayleigh test was performed on a mouse by mouse basis and for the aggregate data. P-values were Bonferroni corrected for multiple comparisons, unless otherwise stated.

To determine the statistical significance of SFC measures, the MI for SFC was compared to an MI distribution obtained using surrogate spiking data generated by using a Poisson process at each time point, using a student’s t-test. The Poisson firing rate was determined by calculating the firing rate of each unit in a 200ms sliding window shifted by 1ms.

## Supporting information

Awake mouse

Fast wave over cortex

Slow wave over cortex

Both fast and slow wave over cortex

## 2 Author Contributions

Conceptualization: AA, DC, MBK, and AP

Data curation: AA, HC, DC, and AP Formal analysis: AA, CB, JL, and AP

Funding acquisition: AA, DC, MBK, and AP

Writing – original draft: AA, MBK, and AP

Writing – review & editing: AA, CB, JL, MBK, DC, and AP

## 3 Acknowledgments

We wish to thank Brenna Shortal for setting up Kilosort for spike sorting. We also want to thank Dr. Andrew Hudson, Dr. George Mashour, Dr. Joseph Cichon, Dr. Andrew McKinstry-Wu, Andrzej Z. Wasilczuk, and Ethan Blackwood for helpful discussions. This research was supported through the Translational Neuroscience Initiative from the Penn Medicine Translational Neuroscience Center (PMTNC), RO1 GM088156 (MBK), RO1 GM124023 (AP), T32 EY007035 (AA), F30 EY029931-01A1 (AA).

## 4 Supplementary Material

– Supplementary figures attached separately
– Video 1: Awake mouse
– Video 2: Average visual evoked fast wave
– Video 3: Average visual evoked slow wave
– Video 4: Both visual evoked waves superimposed

## 6 Data Availability Statement

The raw data and MATLAB code presented in this manuscript will be made available upon request.

