## supplementary figures for "Visual evoked feedforward-feedback travelling waves organize neural activity across the cortical hierarchy in mice"

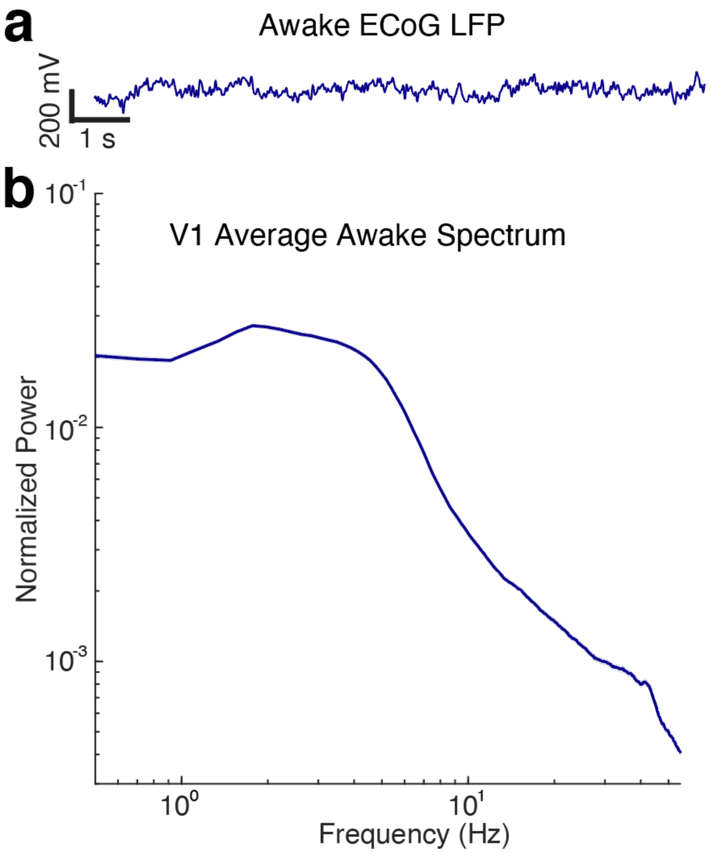


**Supplementary Figure 1: Awake LFP and Power spectrum**

a. Spontaneous LFP recorded over V1 in a representative awake mouse. Note the dominance of high frequency, low amplitude activity
b. Power spectrum of V1 LFP averaged over animals.

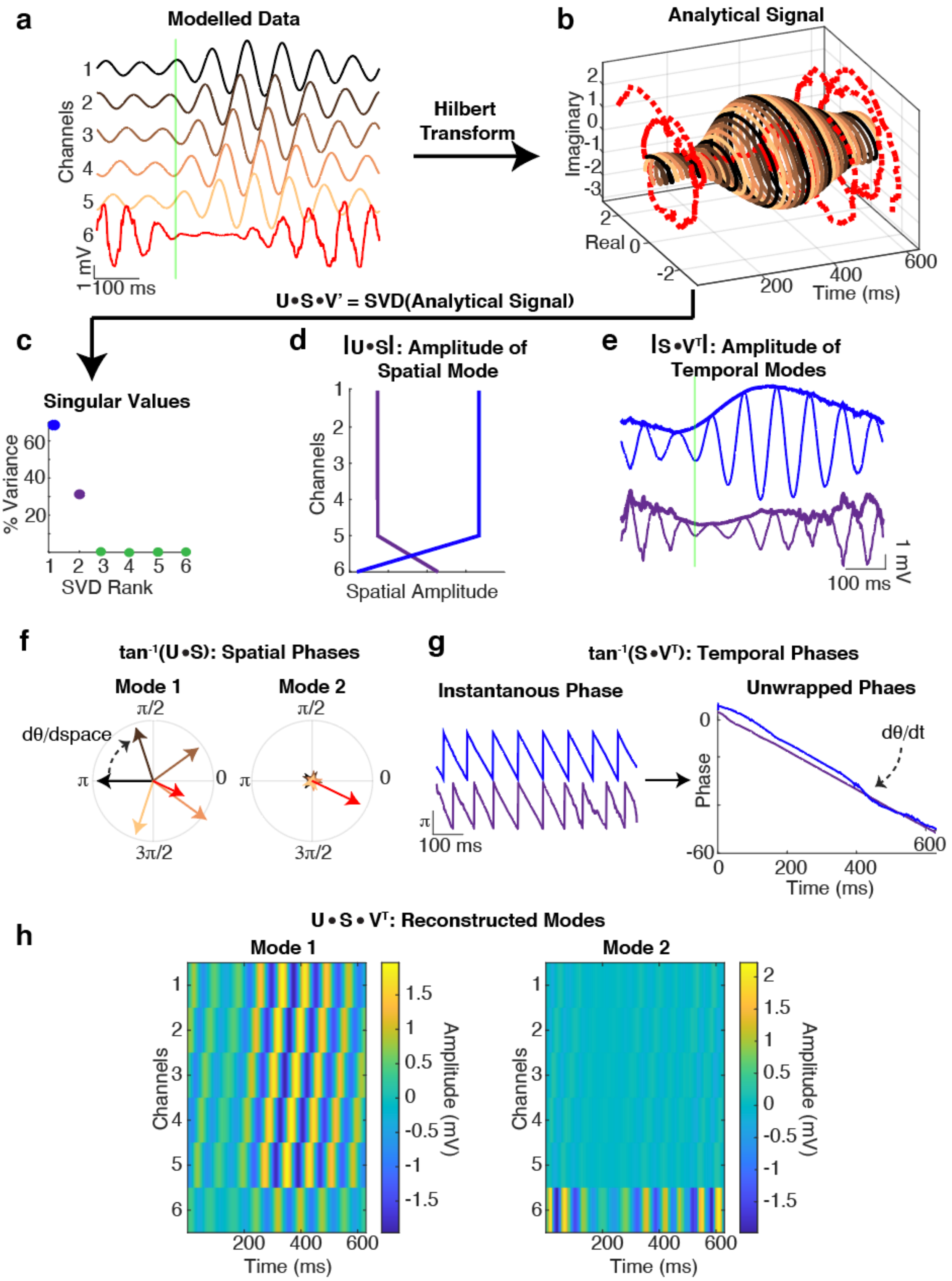
a
Modelled Data
Channels
1 mV
100 ms
Hilbert Transform
b
Analytical Signal
Imaginary
Real
Time (ms)
U•S•V' = SVD(Analytical Signal)
c
Singular Values
% Variance
SVD Rank
d
|U•S|: Amplitude of Spatial Mode
Channels
Spatial Amplitude
e
|S•V^T|: Amplitude of Temporal Modes
1 mV
100 ms
f
tan^-1(U•S): Spatial Phases
Mode 1
Mode 2
dθ/dspace
g
tan^-1(S•V^T): Temporal Phases
Instantanous Phase
Unwrapped Phaes
π
100 ms
Phase
dθ/dt
Time (ms)
h
U•S•V^T: Reconstructed Modes
Mode 1
Mode 2
Channels
Time (ms)
Amplitude (mV)

**Supplementary Figure 2: Singular value decomposition (SVD) of the analytical signal data identifies coherent spatiotemporal activity modes.**

a. Modelled oscillatory activity for 6 electrodes. The signals in the first five electrodes record a traveling wave that propagates from electrode 5 to 1 (black through orange) The sixth electrode (red), is constructed to be independent from others. The green vertical line denotes the stimulus.
b. Analytical (complex valued) signal is obtained using the Hilbert transform from data in A. In the complex plane, the travelling wave appears as phase shifted copies of the same signal and the divergent activity of the sixth electrode is distinct.
c. Singular value decomposition (SVD) can be used to parse the analytical signal into mutually orthogonal modes. Real valued diagonal matrix S encodes the relative contribution of each mode to the overall activity pattern. Complex-valued U and V matrices encode the spatial and temporal characteristics of each mode respectively. SVD identifies only two modes with nonzero singular values, corresponding to the two oscillatory patterns within the signal.
d. The columns of matrix $|\boldsymbol{U}*\boldsymbol{S}|$ encode the spatial amplitude of each mode, and measures how much each electrode contributes to each of the temporal modes. SVD correctly identifies that the first 5 electrodes contribute to the first mode (the traveling wave) equally, whereas mode 2 exclusively involves electrode 6.
e. The temporal amplitudes, computed as $|\boldsymbol{S}*\boldsymbol{V}^{\boldsymbol{T}}|$, reveal the envelope of each mode. Visually responsive modes were defined as modes that increase in temporal amplitude after the stimulus. In this example, only the first mode is visually responsive.
f. The spatial phases are calculated as arctangent of $\boldsymbol{U}$ scaled by the corresponding singular value. The spatial frequency is computed from the phase gradient ($^{d\theta}/_{ds} = F_s$). Here, the phase map of the first mode illustrates that the first five electrodes contribute (black through orange) equally and have a constant phase offset from one another, whereas the sixth electrode (red) has a smaller magnitude. In the second mode, only the sixth electrode has a large magnitude.
g. The temporal phases are calculated as arctan of $\boldsymbol{V}^{\boldsymbol{T}}$. The time derivative of this phase ($^{d\theta}/_{dt} = F_t$) defines the temporal frequency of the mode and is approximated by measuring the slope of the unwrapped temporal phase. The propagation velocity of the mode is then computed as $^{F_t}/_{F_s}$.
h. The activity corresponding to the i-th spatiotemporal mode can be reconstructed as $U_{:,i} * S_{i,i} * V_{i,:}^{T}$. Here, the first mode shows a traveling wave in the first 5 electrodes. The independent spatiotemporal mode in the sixth electrode is present in the second mode.

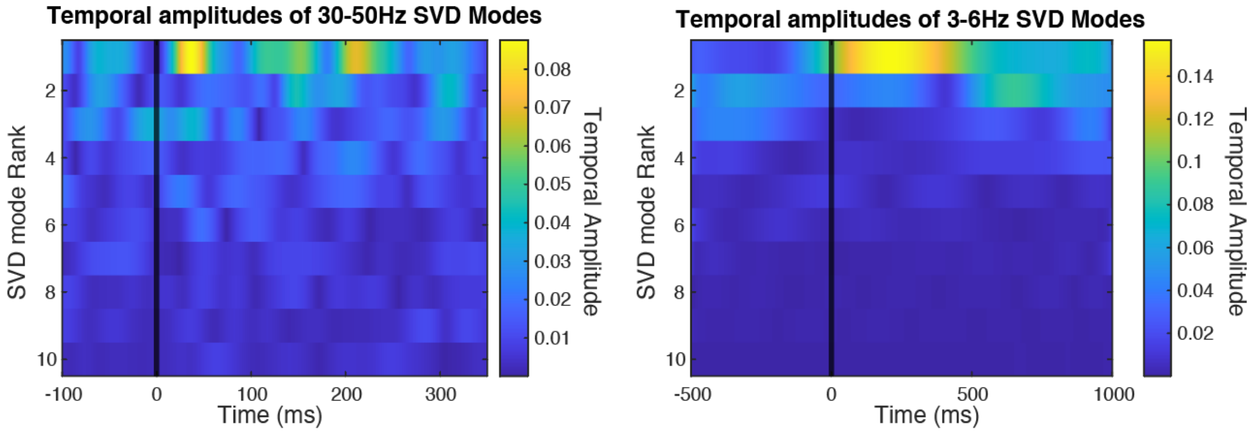


**Supplementary Figure 3: The first mode extracted from SVD typically has the highest post-stimulus temporal amplitude**

Temporal amplitude of each of the first 10 SVD modes of a single trial filtered at 30-50Hz (right) or 3-6Hz (left), by time in ms. The stimulus occurs at the black line. Note in both frequency bands, the first mode has the largest increase in amplitude following the stimulus and is therefore defined as the most responsive visual mode.

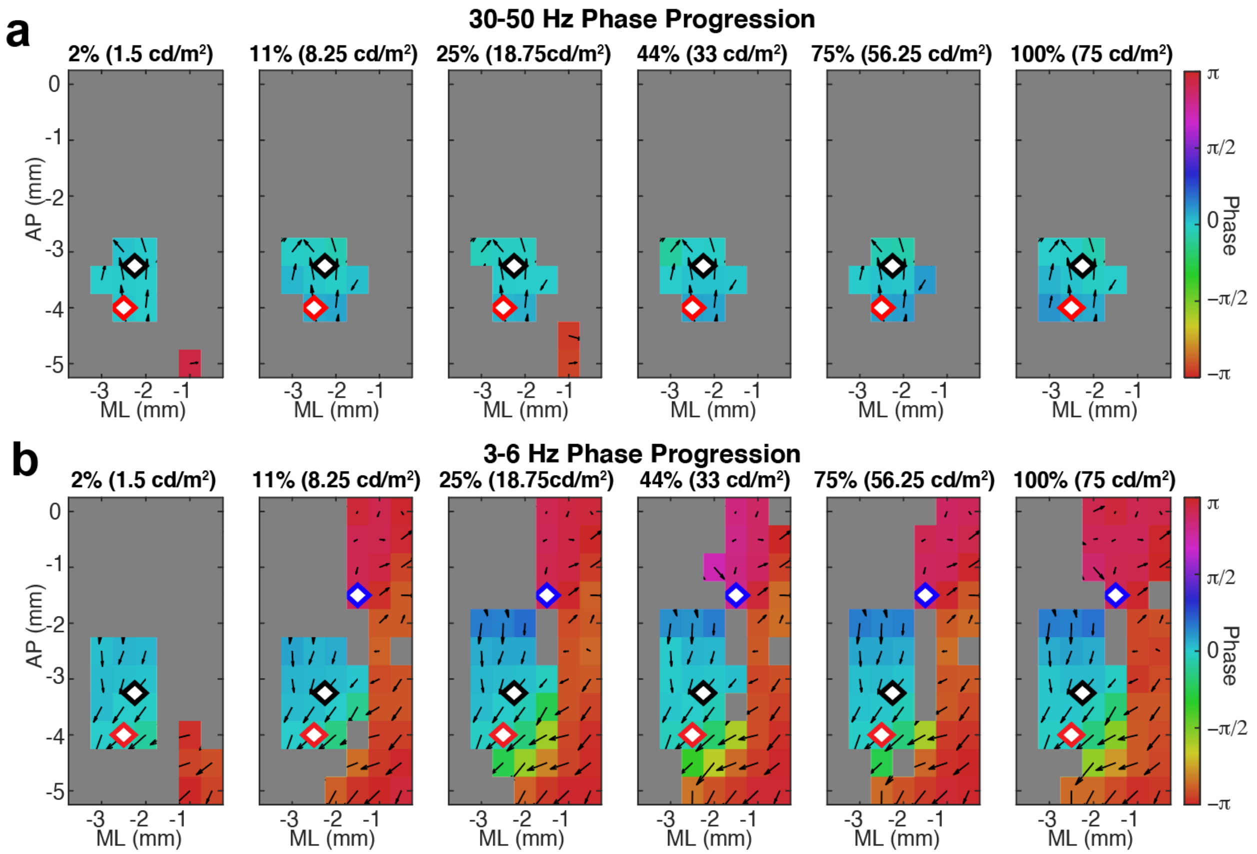


**Supplementary Figure 4: The consistency of the wave propagation pattern depends on stimulus intensity**

a. At each stereotaxic location, the average (over mice) phase offset of the 30-50Hz spatial mode relative to V1 (the black diamond) is plotted in color for each screen luminance, listed as a percent of maximum screen luminance. The red diamond is a different location in V1. Spatial phase gradient is depicted by black arrows. The direction of the arrows shows the direction of spatial phase gradient over trials and mice. The magnitude of the arrows corresponds to the consistency of the angle of the spatial phase gradient over trials and animals. Locations that are grayed out did not meet Bonferroni corrected statistical significance (p-value < 0.0006) across animals.

b. Same plots as in A but for waves identified from 3-6Hz filtered data. The blue diamond denotes the location of the PPA.

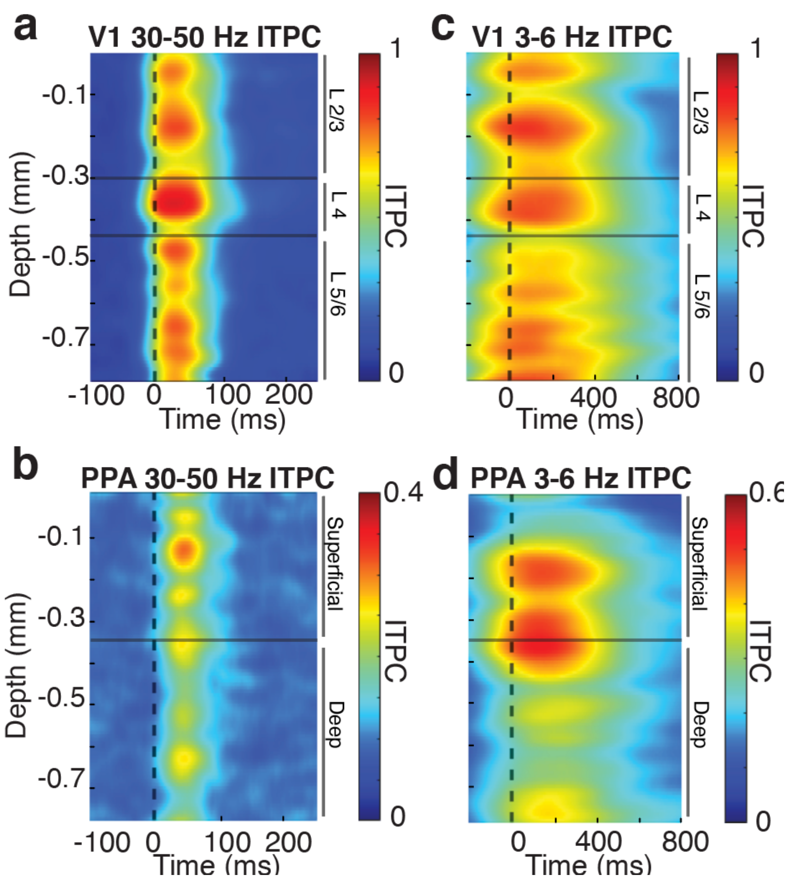


**Supplementary Figure 5: Superficial cortical layers of V1 and PPA contain high ITPC**

a. ITPC of CSD at 30-50Hz as a function of time and depth in V1, averaged over animals. The horizontal lines indicate the supra-, granular, and infragranular layers.
b. Same plots as in A but for PPA. Note that high frequency ITPC is most prominent in superficial cortical layers.
c. ITPC of CSD at 3-6Hz as a function of time and depth in V1, averaged over animals. The horizontal lines indicate the superficial and deep layers.
d. Same plots as C but for PPA. In contrast to V1, ITPC is most dominant in the superficial layers.

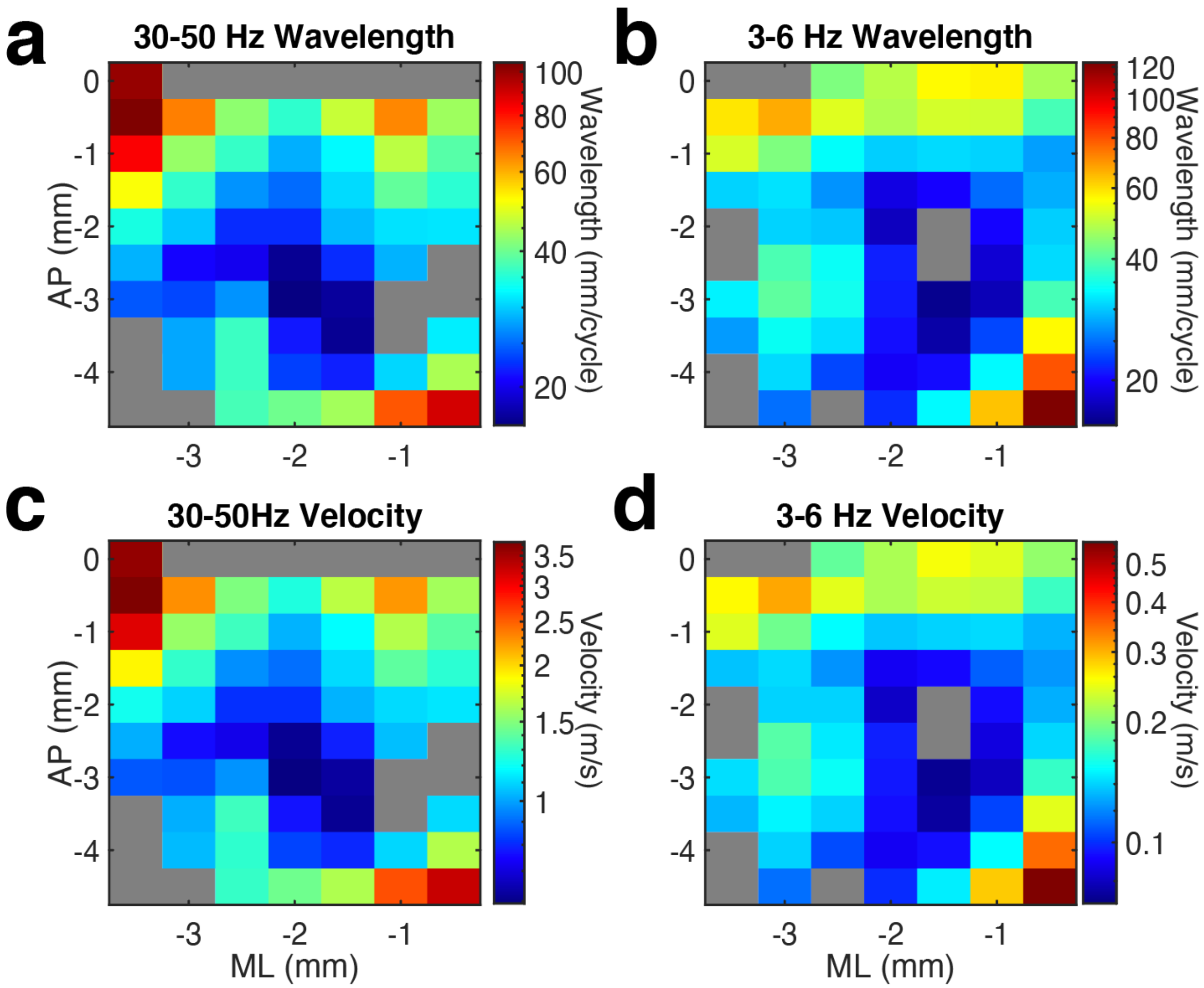


**Supplementary Figure 6: Spatial wavelength and wave velocity are not uniform over space**

a. Spatial wavelength of fast 30-50Hz most visually responsive SVD modes.
b. Spatial wavelength of fast 30-50Hz most visually responsive SVD modes.
c. Velocity of fast 30-50Hz most visually responsive SVD modes.
d. Velocity of fast 30-50Hz most visually responsive SVD modes.

*Note that in all plots, the color axis is in log scale and locations that are grayed out did not meet Bonferroni corrected statistical significance (p-value < 0.0006) across animals.

### Supplementary Videos Legends

**Supplementary Video 1: Awake mouse**
Three seconds of an awake mouse in recording both.

**Supplementary Video 2: Average visual evoked fast wave**
Average of single trial LFP filtered at fast, 30-50Hz, frequencies over the cortical surface during the first 100 ms of the visual evoked response. The flash occurs at time = 0 ms. Note that the visual evoked gamma activity begins caudally within V1 and propagates rostrally.

**Supplementary Video 3: Average visual evoked slow wave**
Average of single trial LFP filtered at slow, 3-6Hz, frequencies, over the cortical surface during the first 600 ms of the visual evoked response. The flash occurs at time = 0 ms. Note that the visual evoked gamma activity begins rostral to V1 in higher order cortical areas and propagates caudally.

**Supplementary Video 4: Both visual evoked waves superimposed**
Superimposition of the average fast and slow filtered waves (amplitude of the signals is normalized to highlight temporal relationships between oscillations). The flash occurs at time = 0 ms. The fast wave begins caudal to the slow wave and travels anterolaterally towards the slow wave initiation zone.
